# High Competition and Selective Extinction: How Biotic and Abiotic Drivers Shaped Speciation and Extinction Regimes in Carnivora

**DOI:** 10.1101/2025.09.22.677728

**Authors:** Lucas M. V. Porto, Tiago B. Quental

## Abstract

Understanding the drivers of biodiversity over time is a central goal in macroevolution. Yet, the relative contributions of biotic and abiotic mechanisms remain unclear, especially at broader phylogenetic and spatial scales. This study investigates how biotic (self-diversity dependence and competition proxies) and abiotic (temperature) factors shaped Carnivora diversification across North America and Eurasia over the last 45 million years. Using a Bayesian framework, curated fossil data, and an expanded method to assess competition intensity at multiple spatial scales, we quantify speciation, extinction, and diversity patterns across 17 families. Our results show that competition significantly influences diversification on both continents. Mechanisms vary by scale, with contrasting associations between diversification rates and predictive timeseries. While competition can hinder speciation by saturating ecological niches, it may also foster diversity via character displacement and niche partitioning, especially under local spatial coexistence. At regional scales, abiotic factors—particularly cooling temperatures and habitat shifts—act as selective extinction drivers, disproportionately affecting specific regions of the morphospace and creating gaps. By integrating temporal and spatial perspectives, our study enhances understanding of how biotic interactions and environmental changes jointly shape biodiversity through deep time.

## Introduction

Understanding the underlying processes responsible for the wax and wane of biodiversity over time is a fundamental goal in macroevolution (Sepkoski 1978; Rosenzweig 1995). The traditional debate primarily focused on the roles of abiotic and biotic factors in shaping diversity dynamics. Studies investigating long-term species diversity changes typically highlighted the predominant role of abiotic factors in driving biodiversity in deep time (Barnosky 2001; Alroy et al. 2008; Benton 2009; Benson et al. 2022). However, there is growing recognition that biotic interactions also play a critical role in modulating diversity patterns over geological time (Ezard et al. 2011; Pires et al. 2015; Cantalapiedra et al. 2018; Close et al. 2019; Hembry and Weber 2020a; Liow and Quental 2025a).

This growing focus on biotic interactions is closely tied to a longstanding debate in paleontology and evolutionary biology: whether species diversity is inherently limited or not at global and regional scales, both among paleontologists (Raup 1972; Sepkoski 1984a, 1978, 1979; Walker and Valentine 1984; Alroy 1996, 2008, 2010; Stanley 2007; Benton and Emerson 2007; Erwin 2008; Quental and Marshall 2013; Marshall and Quental 2016a), as well as neontologists (Harmon and Harrison 2015; Rabosky and Hurlbert 2015). While some studies argue that diversification is unconstrained (e.g. Benton and Emerson 2007; Harmon and Harrison 2015), others suggest that diversification slows down over time due to diversity-dependent diversification (Sepkoski 1984b; Rabosky and Hurlbert 2015), which might include or not a fixed carrying capacity (Rabosky 2013a; Marshall and Quental 2016b; Foote 2023). In this view of limited growth of biodiversity, an increase in diversity within a clade should limit speciation or increase extinction risk (Sepkoski 1996; Rabosky 2013b; Rabosky et al. 2014; Ezard and Purvis 2016). In such diversity dependence scenario the drop in speciation would reflect saturated ecological niches (Simpson 1953; Sepkoski 1996; Schluter D 2000), while and increase in diversity would lead to a higher risk of extinction of ecologically similar species reflecting competitive exclusion (Van Valen 1985; Sepkoski 1996).

Most studies have used diversity-dependence as a proxy for interspecific competition, either within or between clades (e.g. Sepkoski et al. 2000; Silvestro et al. 2015a; Pires et al. 2017a), assuming that higher diversity inherently intensifies competition. As explicitly pointed out by Foote (2023) “…*prevalent biotic interactions need not imply diversity dependence, and diversity dependence need not imply logistic or equilibrial diversity with a fixed carrying capacity*.” Hence although the assumption that diversity-dependence represents biotic interactions might be often reasonable, it might oversimplify the complex interplay of factors that determine the fate of lineages. For example, Guo et al. (2023) suggested that the strong negative correlation between bivalve diversity and brachiopod origination rates in the fossil record, often attributed to competition, may instead result from post-extinction rebounds in origination when diversity is low, rather than direct evidence of competition (but see Liow et al. (2015) for a counter argument).

The role of biotic interaction on driving the fate of different populations and microevolution is well accepted among ecologists and evolutionary biologists, but scaling them up to understand the dynamics at higher levels of diversity remains challenging (Jablonski 2008; Hembry and Weber 2020b). Moreover, accessing the potential role of biotic interactions in deep time is difficult due to the historical nature of the investigation. Even though the fossil record is the more direct way to document the origin and extinction of different lineages, their incompletness (Raup 1979), and the difficulty of directly observe species interactions (but see Lidgard et al. (1993) and Liow et al. (2016) for direct evidence of interspecific competition in the fossil record), means that the effect of biotic interactions have relied on rather indirect evidence, most notably diversity dependence.

In this study, we aim to assess the potential role of biotic and abiotic factors on controlling the diversification dynamics of Carnivora across North America and Eurasia. The order Carnivora provides an ideal model for addressing these questions. First, it presents a fossil record spanning over 45 million years and well-documented taxonomic diversity (Werdelin and Turner 1996; Van Valkenburgh 1999; Wang and Tedford 2008). Second, their eco-morphological diversity—ranging from small hypocarnivorous plant consumers to large hypercarnivorous hunters—is well characterized (Van Valkenburgh 1999, 2007; Balisi et al. 2018), making them a robust system for investigating the interplay of biotic and abiotic factors in shaping diversity dynamics. Third, previous studies on Carnivora have suggested a significant role of competition (here measured by diversity dependence) in shaping diversification dynamics (Silvestro et al. 2015b; Pires et al. 2017b).

To more directly evaluate the role of interspecific competition we used the approach recently developed by Graciotti et al. (2025) also discussed in Liow and Quental (2025b). This approach uses a framework that considers both temporal coexistence and spatial and ecological overlap. Therefore, it represents an attempt to characterize the intensity of interspecific competition by measuring how ecologically similar species are, and how they coexist in time and space. We used a Bayesian framework (PyRate) (Silvestro et al. 2014) to estimate the rates of speciation and extinction and their potential association with different time series representing either biotic or abiotic factors. This was done by applying a multivariate birth-death model (PyRateMBD) (Lehtonen et al. 2017a) across Carnivora in North America and Eurasia. We also included diversity as a surrogate for biotic interactions, allowing us to compare the inference that has been traditionally used to our new proxy for interspecific competition, a measure of ecological similarity while taking into account the temporal and spatial overlap of species.

## Methods

### Data Collection and Preprocessing

Fossil occurrence data of Carnivora was manually downloaded from the Paleobiology Database (PBDB) on 06/07/2022 and from the New and Old Worlds fossil mammal database (NOW) on 01/07/2022. Occurrences that were not identified to the species level, or showed some taxonomic uncertainty, were removed from the dataset. Aquatic species, based on “taxonomy” (all species that belong to aquatic families), were removed latter from our dataset.

Initially, the NOW database contained 8,039 occurrences, while the PBDB dataset included 6,949 occurrences. To improve occurrence temporal resolution, we filtered out any occurrences with stratigraphic intervals longer than 6 million years (Figure S1). This was done to prevent overestimating species true times of speciation and extinction in the following Bayesian framework.

To eliminate potential duplicated occurrences when merging the PBDB and NOW datasets, we implemented a custom code (Appendix) that identified and removed duplicate records based on specific criteria. Before removing duplicates, we proceed to harmonize the taxonomic names. This was done by comparing the species in our datasets with those in the CarniFOSS dataset from Faurby et al. (2021b). CarniFOSS provides a curated list of Carnivora species, with corrections for potential typographical errors and synonyms. Through this comparison, we identified and removed 62 species (and their occurrence records) from our dataset that were either incorrect or inconsistent with the CarniFOSS reference (Appendix - Tables S1 and S2).

To find duplicated records the code searched for occurrences with the same generic and specific epithets, a spatial difference of less than one degree of latitude and longitude, and a temporal difference of less than six million years. If two occurrences—one from NOW and one from PBDB—met these criteria, the occurrence from NOW was excluded as a duplicate. This approach allowed us to refine the dataset by removing redundant data, thereby enhancing the reliability of our analysis (Figure S2). After removing duplicates, we ended up with an unbalanced distribution of fossil records over Africa, Europe, N. America, and S. America (Figure S3). We decided to focus our analysis on the geographical regions with more data. Thus, records from Africa and S. America were removed. Following these cleaning steps, our initial pool of 1,155 unique species (combining NOW and PBDB data) was reduced to 798 species. The final dataset retained 7420 records (Appendix – Table S3), ensuring a robust and accurate representation of Carnivora diversity for subsequent analyses.

From PBDB and NOW we also retrieved information regarding the geographical location of fossil occurrences, as each occurrence is identified with an “ID” number that associates it with a specific locality from where each specimen was retrieved. This information was used to group multiple occurrences that were found together into a “local assemblage”. Each occurrence is also georeferenced with latitude and longitude coordinates. Such geographical information allowed us to reconstruct some aspects of species distribution in the fossil record. At the end of the cleaning steps, we had seventeen clades, which could be present in both continents (North American and Eurasian) or only in one (Figures S4 and Table S3). All cleaning steps related to data collection and data cleaning were performed using the R software (v. 4.3.2) (R Development Core Team 2023).

### Ecomorphological Data

Although the framework utilized here, adapted from Graciotti et al. (2025), allows species to be described across various dimensions of morphospace, due to lack of data, in our study, we focused only on body size. Body size is commonly recognized as a robust variable for describing species ecology, as it correlates with numerous aspects of morphology, physiology, and resource utilization (Van Valkenburgh et al. 2004; Bonner 2006; Balisi et al. 2018). We utilized the published CarniFOSS dataset for body size data on Carnivora (Faurby et al. 2021a) (Appendix – Table S5). Out of the 798 species from which we had fossil occurrence data, 16 did not have body mass information in CarniFOSS. Those were removed from the analysis.

### Initial PyRate Analysis

We employed PyRate, a hierarchical Bayesian framework that estimates preservation and diversification processes while explicitly addressing the incompleteness of the fossil record using all known occurrences of a lineage (Silvestro et al. 2014, 2019). We used all fossil occurrences, separated into 2 different continental regions (North America lineages and Eurasian Lineages), to estimate species “true” origination and extinction times. We configured our analyses using the mG + qShift model, running 30.000.000 RJBDMCMC iterations, sampling every 10.000 iterations, and discarding the first 10% as burn-in to obtain posterior parameter estimates. These settings allowed us to model preservation processes that account for variability over time (Time-variable Poisson Process - TPP) and across lineages, a more realistic method to describe fluctuating fossil preservation patterns (mG + qShift model in PyRate). To evaluate the possibility of temporal variation in preservation we grouped stages to define different time windows. The boundaries for these windows were defined based on how the fossil records of each group were distributed in time. We opted to use intervals that minimized the percentage of occurrences that crossed such boundaries. For North America, the intervals were set at 0.2, 4.9, 13.68, 24.8 and 41 million years ago (Figure S5), and for Eurasia we set the intervals as 2.58, 14.2, 23.03 and 38 million years ago (Figure S6). We utilized such times frames to accommodate a roughly homogeneous number of occurrences among intervals across the whole-time frame, but also to minimize the number of occurrences that would cross intervals. To account for uncertainty in fossil ages, we randomly sampled ages within occurrence timespans, generating 20 temporal replicates, and conducted PyRate analyses on each replicated dataset. From these analyses, we derived the general trends in speciation and extinction rates, as well as the times of origin and extinction for each species, which were then used in the subsequent steps to define temporal coexistence.

### Temporal and Spatial Coexistence

Temporal coexistence was estimated using the estimates of the Time of speciation and Time of Extinction (TS and TE) derived from the 20 PyRate replicates. Longevity for each species was calculated from the mean value derived from posterior distributions of speciation and extinction times. The temporal species coexistence was measured in 0.1 million-year intervals for each of the 20 replicates, producing 20 time series, which could represent an underestimate of diversity (the number of sampled species at each point in time), but were in fact used to calculate our disparity metrics (see description below). To actually estimate the number of species we used the mcmcDivE approach implemented in PyRate which takes into account preservation estimates and implements a hierarchical Bayesian model to estimate diversity that explicitly models the non-sampled species, generating a “corrected” diversity curve.

To take into account spatial coexistence, we used two approaches that use the spatial location of fossil occurrence data. Our conservative “Site” method only considered species found in the same locality as coexisting. In our more permissive “Regional” method we considered all species within North America or Eurasia to be potential competitors. In the “Site” scenario, we used the “collection” identifier from PBDB and NOW datasets to generate one continuum ID number for each fossil site. Species in the same fossil site ID number were considered to coexist in this metric.

### Competition Intensity Framework

To better capture competition dynamics, we used a recently developed framework developed by Graciotti et al. (2025). Such a method allowed us to measure the intensity of competition within both North American and Eurasian lineages of Carnivora by assuming that eco-morphological distance is inversely proportional to the intensity of competition. The closer species are in an eco-morpho-space the stronger the intensity of competition. Here we used body size as our morphospace variable, and distances were estimated using the Mean Nearest Neighbor distance (MNND). The MNND was estimated for several time points spaces by 0.1 million-year intervals. Only species that coexisted in time and space at each time point were included. This produced two time series, one for each spatial metric (“Site” and “Regional”). Such a time series were used as input for the correlation analyses described in the next section.

In the MNND metric, only the distances between each species and its nearest neighbor in the morphospace are calculated (Foote 1990; Guillerme et al. 2020). Since only the closest distances are taken into account, MNND excludes potential competition from species that are less likely to compete. We expect MNND to reflect competition from the most direct competitors. As the MNND values increase, species are getting further away from each other in the morphospace; hence, we anticipate a decrease in competition intensity over time.

### Pyrate MBD (Multivariate Birth-Death model)

We utilized the multivariate birth-death (MBD) model within the PyRate software (Lehtonen et al. 2017b), to evaluate how both biotic and abiotic factors influenced the diversification patterns of Carnivora. In this model, speciation and extinction rates are connected to time-continuous variables through an exponential function, where the observed rates are determined by the combination of baseline rates, correlation parameters, and the provided time series for the different variables being used. PyRateMBD uses an MCMC algorithm to estimate the correlation parameters that each variable has with speciation and extinction rates (Gλ and Gµ). A positive value of G indicates a positive correlation between the factor and the rate, while a negative value suggests the opposite.

We applied the multivariate analysis to five different time-series: Regional and site diversity, regional and site MNND, and temperature. Temperature data was obtained from the R package RPANDA (v. 2.4) (Morlon et al. 2016) and represents estimates of global temperature inferred from O18. Correlations were performed separately for North American and Eurasian lineages. Prior to running the analysis, all time-series were rescaled between 0 and 1. The MBD model was run for 30 million generations, with 1000 posterior samples saved. For each TS and TE dataset, from the initial PyRate analysis, we performed 20 independent replicates (20 independent PyRateMBD analysis), and the results from these replicates were combined after discarding the first 10% as burn-in. We assessed convergence using Tracer (v.1.7.1) (Rambaut et al. 2018), and for any replicates with ESSs < 200, additional generations were analyzed. We then calculated the 95% highest posterior density intervals for the correlation parameters, along with the average shrinkage weights. Following Guo et al. (2023) and Jouault et al. (2024), relationships were considered significant if the shrinkage weight for a correlation was equal or higher than 0.5.

To assess if diversification drivers may change over time, we divided the whole-time interval studied here into four time windows: window 1 (35 - 31.5 Ma), window 2 (31.5 to 23.5 Ma), window 3 (23.5 to 7 Ma), and window 4 (7 Ma to the present). Analysis for North America and Eurasia were set with the same time windows for comparison. The selection of the four-time windows was based on a visual inspection of the diversification rates for both clades, along with their diversity curves (Figures S7 and S8). We aimed to identify periods where phases of radiation, decline, and equilibrium—judging from both the diversification rates and diversity trajectories—were distinct, allowing us to separate these moments for more accurate correlation analyses.

All R and PyRate scripts used during our analysis are available as supporting information (Appendix) together with a “roadmap” to guide over each step.

## Results

### Diversification Dynamics

Our results reveal marked temporal variation in speciation, extinction, and net diversification rates for Carnivora in North America and Eurasia, producing dynamic patterns (Figures 1 and 2). In North America, speciation peaked before 35 Ma, then declined sharply and remained low, with brief increases around 16 and 5 Ma (Figure 1B). Extinction stayed low until ∼3 Ma, with a minor peak at ∼13 Ma, then rose steadily (Figure 1C). Net diversification was initially high before 35 Ma (Figure 1A), followed by a prolonged near-zero phase, interrupted by two small positive pulses and a slight negative dip at ∼13 Ma. The recent rise in extinction, combined with low speciation, led to a pronounced negative diversification phase from ∼3 Ma to the present.

**Figure 1.**
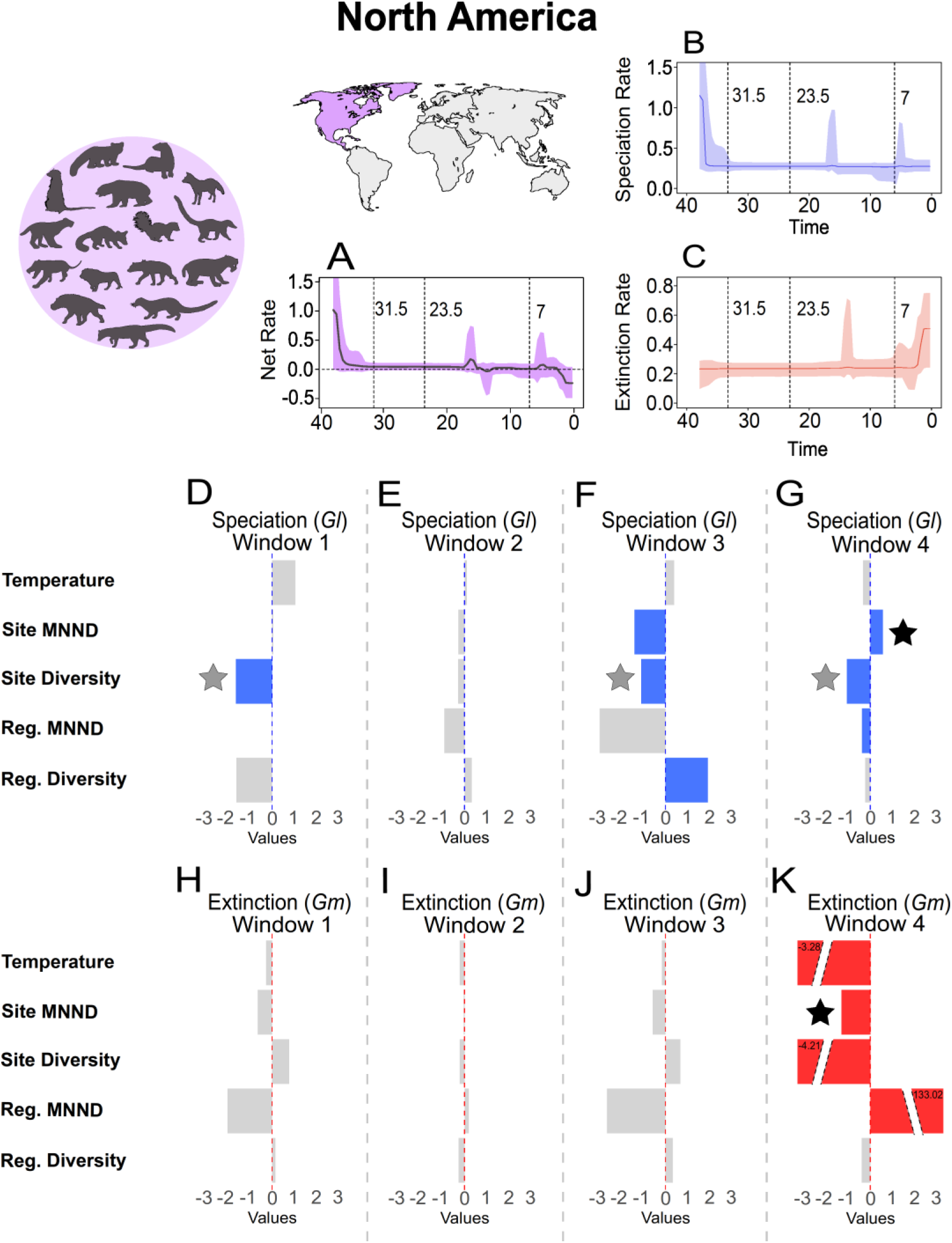
Net diversification (A), speciation (B), and extinction (C) rates of Carnivora in North America, with correlations to abiotic and biotic factors across four temporal windows. Panels D–G show correlations with speciation rates, and H–K with extinction rates, for the five timeseries analyzed (temperature, site MNND, site diversity, regional MNND, and regional diversity). Colored bars (blue: speciation, red: extinction) indicate significant correlations (shrinkage weight > 0.5), while grey bars indicate non-significance. Temporal windows: 1 (35–31.5 Ma), 2 (31.5–23.5 Ma), 3 (23.5–7 Ma), 4 (7 Ma–present). The dashed line within each temporal window (D–K) represents zero correlation; bars left of it indicate negative correlations, and bars right indicate positive correlations. The “star” symbol at the bars represents correlations that we would expect under a model of interspecific competition, where black stars represent scenarios that more directly measure niche saturation, and grey stars represent scenarios that niche saturation is measured indirectly by diversity dependence.

**Figure 2.**
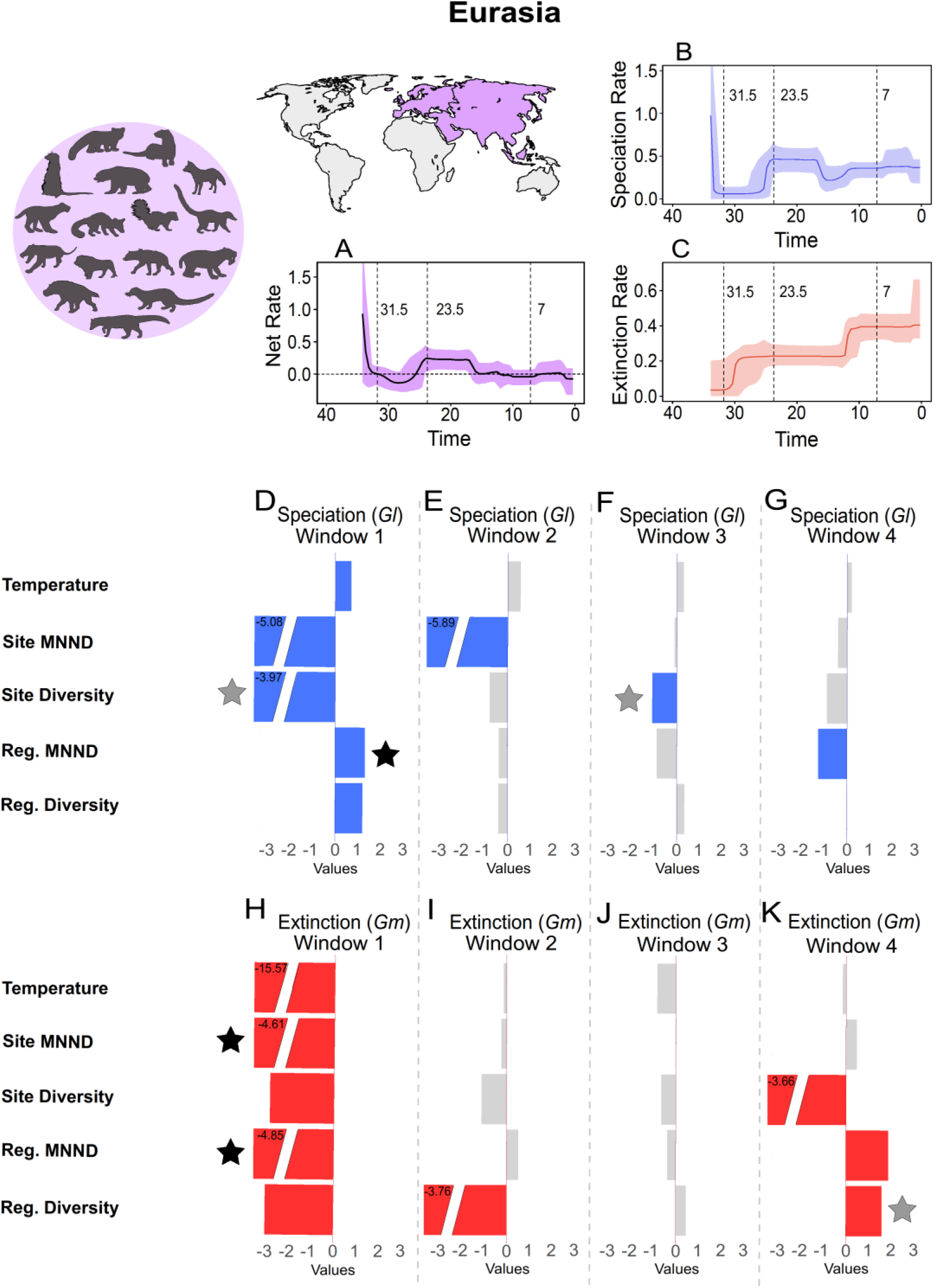
Net diversification (A), speciation (B), and extinction (C) rates of Carnivora in Eurasia, with correlations to abiotic and biotic factors across four temporal windows. Panels D–G show correlations with speciation rates, and H– K with extinction rates, for the five timeseries analyzed (temperature, site MNND, site diversity, regional MNND, and regional diversity). Colored bars (blue: speciation, red: extinction) indicate significant correlations (shrinkage weight > 0.5), while grey bars indicate non-significance. Temporal windows: 1 (35–31.5 Ma), 2 (31.5–23.5 Ma), 3 (23.5–7 Ma), 4 (7 Ma–present). The dashed line within each temporal window (D–K) represents zero correlation; bars left of it indicate negative correlations, and bars right indicate positive correlations. The “star” symbol at the bars represents correlations that we would expect under a model of interspecific competition, where black stars represent scenarios that more directly measure niche saturation, and grey stars represent scenarios that niche saturation is measured indirectly by diversity dependence.

In Eurasia, speciation peaked earlier (∼33.5 Ma; Figure 2B), declined sharply until ∼26 Ma, then rebounded between 26–16 Ma, stabilizing thereafter. Speciation magnitude was consistently higher than in North America. Extinction showed episodic increases, notably at ∼30 and ∼13 Ma (Figure 2C), but remained stable otherwise. Net diversification peaked at ∼33.5 Ma (Figure 2A), then dropped into negative values due to declining speciation and rising extinction. Between 25 and ∼13 Ma, diversification stabilized positively before declining again, fluctuating near zero over the last 15 Ma.

Eurasian Carnivora exhibited greater temporal fluctuation than North American lineages, though both regions share an early diversification peak, a long equilibrium phase, and a recent decline. Eurasia also experienced a distinct positive diversification phase from the Late Oligocene to mid-Miocene, following a pronounced negative phase around 25–30 Ma. In contrast, North America shows a more subdued pattern, with a prominent negative diversification phase only near the present. These findings underscore both regional differences and shared macroevolutionary trajectories in Carnivora diversification.

### Abiotic and Biotic Time-series

Figures 3 and 4 present five time-series correlating with speciation and extinction rates in North America and Eurasia. In North America, Carnivora regional diversity gradually rises between 38–32 Ma, stabilizes until ∼25 Ma, and peaks around 15 Ma, coinciding with elevated speciation (Figure 3B). A sharp decline follows ∼2 Ma later, likely linked to a temporary extinction spike. Diversity then remains stable until the Holocene, when it drops abruptly. Site-level diversity mirrors these trends, peaking earlier in the Miocene with a gentler mid-Miocene decline. While regional diversity reflects abrupt shifts, site diversity smooths these changes, indicating gradual variation in species coexistence (Figure 3C).

**Figure 3.**
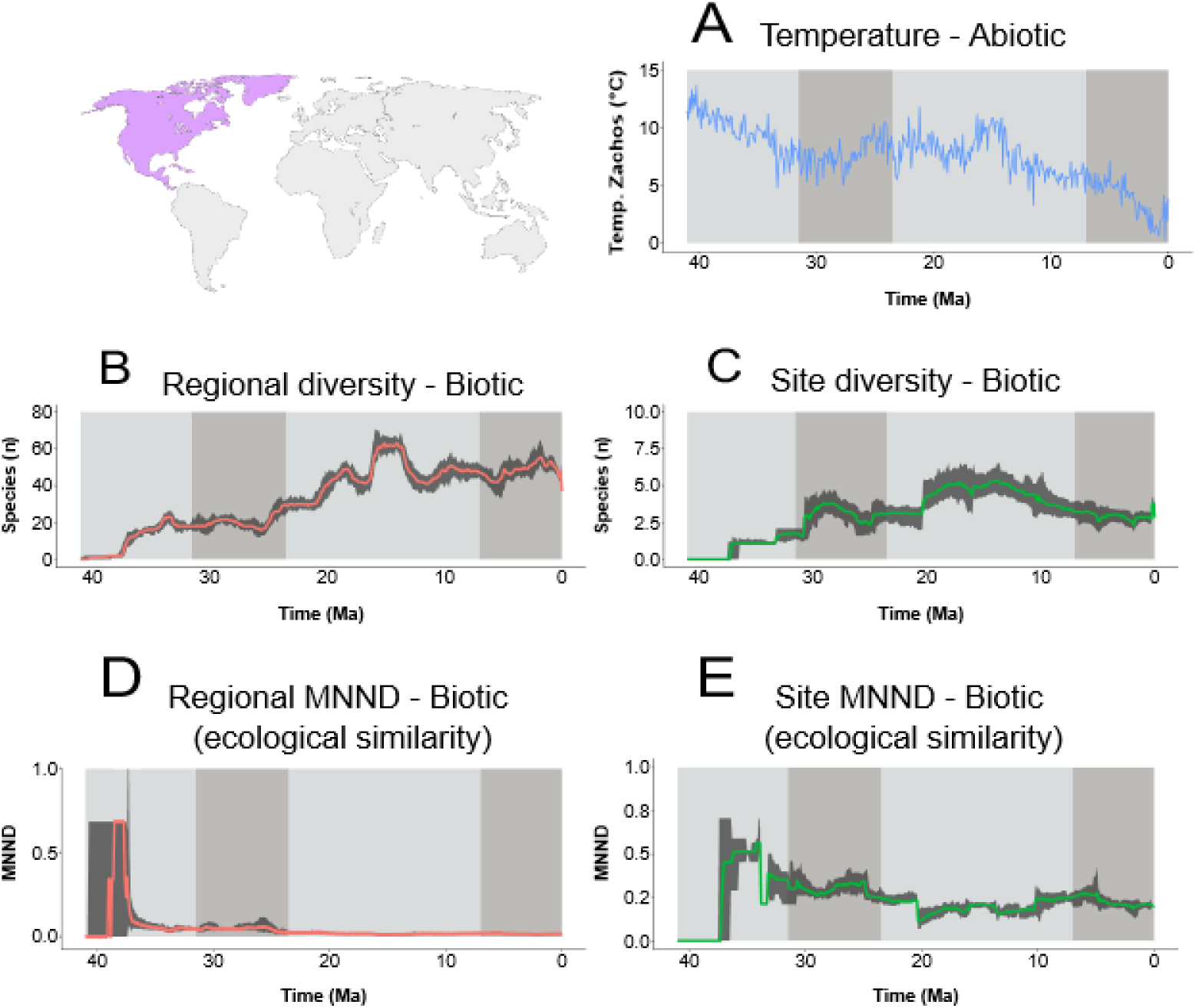
Five timeseries used in the study for Carnivora in North America: (A) Temperature as an abiotic factor. (B) Regional diversity and (C) site diversity as measures of diversity-dependence. (D) Regional MNND and (E) site MNND as proxies for competition intensity (biotic interactions). Solid colored lines represent the median values of each timeseries, while the shaded areas show the range (maximum and minimum values) across different replicates. The gray shaded vertical boxes indicate the temporal windows (used in PyRate MBD analyses), designed to capture key phases in the evolutionary history of Carnivora.

**Figure 4.**
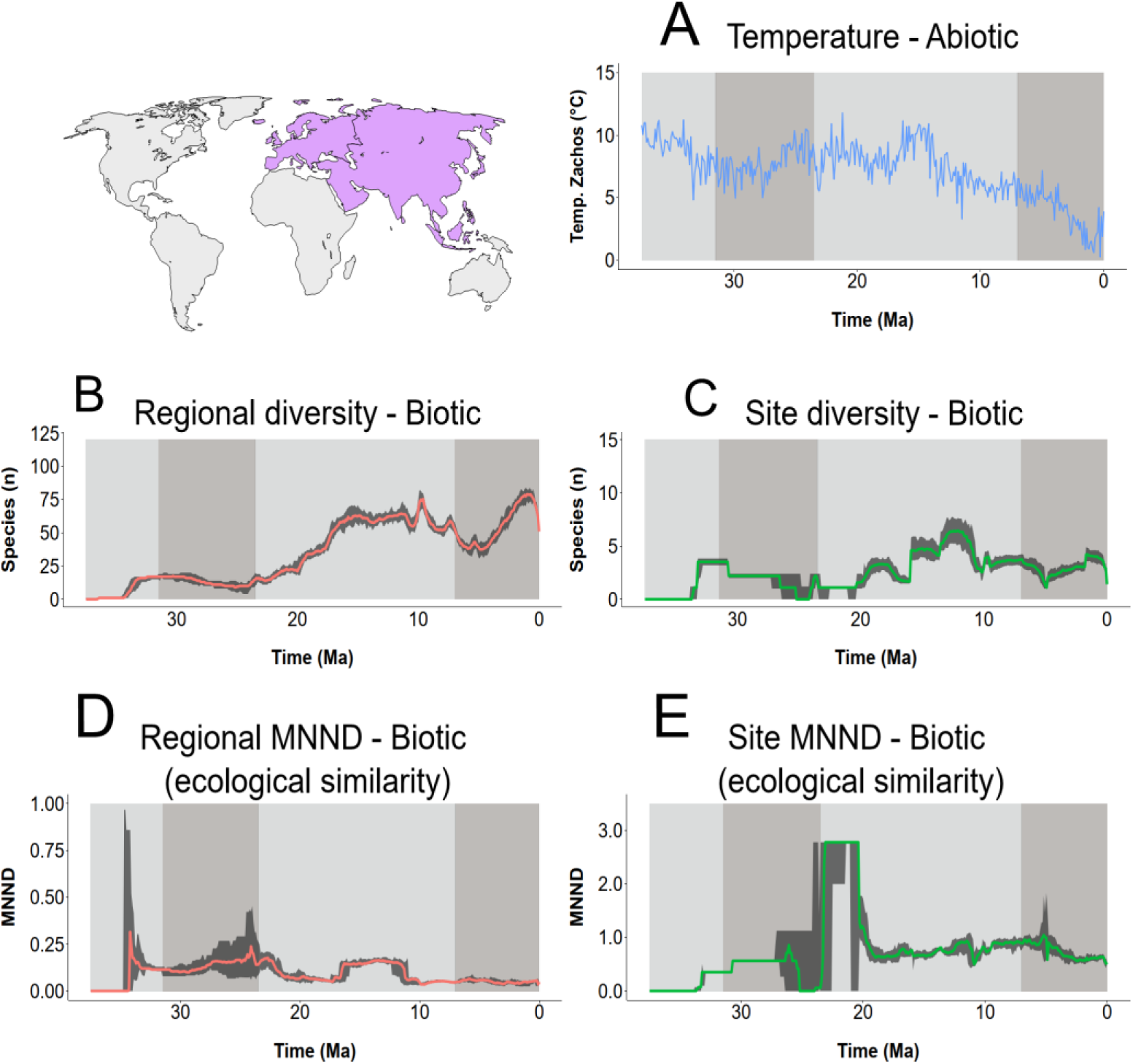
Five timeseries used in the study for Carnivora in Eurasia: (A) Temperature as an abiotic factor. (B) Regional diversity and (C) site diversity as measures of diversity-dependence. (D) Regional MNND and (E) site MNND as proxies for competition intensity (biotic interactions). Solid colored lines represent the median values of each timeseries, while the shaded areas show the range (maximum and minimum values) across different replicates. The gray shaded vertical boxes indicate the temporal windows (used in PyRate MBD analyses), designed to capture key phases in the evolutionary history of Carnivora.

Mean Nearest Neighbor Distance (MNND) at both scales (Figures 3D and 3E) shows early radiation with a sharp increase, followed by decline due to ecological saturation. Regional MNND remains low and stable from the Miocene onward, while site-level MNND fluctuates more, reflecting localized dynamics.

In Eurasia, regional diversity rises quickly to a low-diversity equilibrium (35–25 Ma), then increases steadily during the Miocene. Around 10 Ma, diversity declines due to rising extinction, stabilizes, and rises again in the Pliocene (∼5 Ma) (Figure 4B). Site-level diversity fluctuates after a slight rise at ∼32 Ma, peaking near 12 Ma (Figure 4C).

MNND patterns differ between continents. In Eurasia, regional MNND peaks early and declines gradually (Figure 4D), while site-level MNND peaks later (∼23 Ma), aligning with increased speciation and positive diversification (Figure 4E). Temperature trends (Figures 3A and 4A) show overall cooling from the Eocene, with relative stability and slight warming in the early Miocene.

Species coexistence variance (Figures S9A and S9B) increased during the mid-to-late Miocene in both continents, suggesting greater heterogeneity in species richness. Earlier intervals showed lower, more homogeneous coexistence patterns.

### Association Between Time-series and Macroevolutionary Dynamics

#### North America

All time-series showed significant correlations with Carnivora speciation or extinction rates in at least one of four time windows (Figures 1D–1K). In time window 1 (Figure 1D), site diversity negatively correlates with speciation, suggesting diversity-dependence during early Carnivora radiation in North America. No significant correlations were found in time window 2 (Figure 1E).

In time window 3 (Figure 1F), three variables correlate with speciation: site diversity (negatively), again indicating diversity-dependence; regional diversity (positively), suggesting broader-scale external drivers; and site MNND (negatively), contrary to expectations, implying new species emerged closer in morphospace.

Time window 4 (Figure 1G) reveals more complex dynamics. Site MNND correlates positively with speciation, while site diversity maintains a negative correlation. Regional MNND shows a weak but significant negative correlation.

For extinction rates, no significant correlations appear in windows 1–3 (Figures 1H–1J). In window 4 (Figure 1K), multiple variables show strong correlations: regional MNND correlates positively with extinction, while site MNND correlates negatively, suggesting lower extinction when species are more morphologically dispersed. Site diversity also correlates negatively with extinction, challenging predictions from interspecific competition models. Additionally, temperature shows a negative correlation with extinction, indicating that lower temperatures were associated with higher extinction rates—likely reflecting major environmental shifts that impacted Carnivora diversity in North America over the past 7 million years.

#### Eurasia

In time window 1 (Figures 2D and 2H), all variables significantly correlate with both speciation and extinction rates. Site diversity supports diversity-dependence in speciation but not extinction. Regional diversity behaves unexpectedly for both. Site MNND shows negative correlations with both rates, while regional MNND aligns with competition dynamics—speciation increases and extinction decreases as species become more morphologically distinct. Temperature also plays a role: higher temperatures boost speciation and reduce extinction, with a stronger effect on extinction.

In time window 2 (Figures 2E and 2I), site MNND and regional diversity show significant negative correlations with speciation and extinction, respectively. Other variables show no significant relationships, suggesting distinct ecological processes operating at site versus regional scales.

In time window 3 (Figure 2F), only site diversity significantly correlates with speciation, reinforcing the idea of diversity-dependence at localized sites where species interact directly.

Time window 4 reveals contrasting patterns in Eurasia. Speciation negatively correlates with regional MNND (Figure 2G), suggesting competition promotes lineage formation. Extinction correlates negatively with site diversity but positively with regional diversity (Figure 2K), implying that broader-scale diversity intensifies extinction, consistent with diversity-dependent models. Regional MNND also positively correlates with extinction, indicating that greater ecological distinctiveness increases extinction risk.

Together, these patterns suggest that different ecological mechanisms influence speciation and extinction across spatial scales and time periods.

## Discussion

The diversification dynamics of Carnivora in North America and Eurasia reveal complex interactions shaped by biotic and abiotic factors. Diversity-dependence and distance metrics (both proxies for interspecific competition) more often influenced speciation in North America, while in Eurasia both rates showed similar evidence of interspecific competition. Even though temperature is not often associated with changes in speciation and extinction, unexpected (from the point of view of interspecific competition) associations between the other time series and both rates suggests that external environmental factors might also play an important role. Our findings expand on Graciotti et al. (2025), emphasizing temporal and spatial variations in these processes. Here, we present the contrasting evolutionary dynamics experienced by carnivoran lineages in North America and Eurasia. Below, we explore key findings for each continent and time window.

### North America

In North America, diversity-dependence emerges as a strong factor influencing speciation, particularly when coexistence is measured at the site level, where species directly interact. During the earliest time window, speciation aligns with classic diversity-dependence patterns, where it drops as species start to encounter more species at the site level. This phase corresponds to ecological saturation, as niches are filled, limiting new species’ opportunities to establish themselves. Interestingly, extinction rates during this time window do not display significant signals of diversity dependence. This absence may reflect the nature of a radiation phase, where the expansion into available niches more directly influences speciation over extinction (Givnish 2010; Bouchenak-Khelladi et al. 2015). This pattern could also be explained by ecological speciation, in which new species arise already differentiated enough to minimize interspecific competition. As a result, extinction appears less constrained, as species are formed in ways that reduce competitive overlaps, a process that becomes less feasible as other species are added at the site level. Interestingly, a significant signal of diversity-dependence in speciation is not recovered at the regional level, although the negative direction of association detected is what we would expect under a model of diversity-dependence.

Following this initial radiation, the second temporal window in North America marks a phase of stability in diversification dynamics. During this period, neither speciation nor extinction correlates significantly with diversity metrics. This could suggest a diminished role for both biotic and abiotic factors, but could also reflect the nature of a stable phase where detecting a significant association would be hindered by no variation. We suspect this latter option is the case. Hence this equilibrium phase likely reflects a saturated adaptive zone, where ecological opportunities had been fully exploited. While similar patterns of stability have been documented during the Neogene (Liow and Finarelli 2014), our findings indicate that this equilibrium phase occurred earlier in North America than in Eurasia where a more stable dynamics only happens closer to the present (Figure 2).

By temporal window 3, an intriguing deviation from our competition effects original expectations arises, with regional diversity positively correlating with speciation. We suspect this positive association is related to immigration rather than speciation *per se*, because the method we used does not distinguish immigration from speciation, and any new species is counted as a speciation event. In fact, around that time, there is an influx of Carnivora into North America, mostly driven by immigrant Feliformes from Eurasia (Pires et al. 2015). This immigration likely repopulated the niches left vacant during the “cat-gap” (25– 18.5 Ma), when few cat-like species are observed in North America (Van Valkenburgh 1999). Hence, a positive association between speciation rate and diversity we recover likely reflects this expansion of the morphospace produced by those immigrants. That said, it is also possible that higher regional diversity could also act as an ecological catalyst rather than a limitation for speciation. Simultaneously, site MNND’s presents a negative correlation with speciation, during the time window when MNND time series showed a marked drop around 20 Ma. This suggests that as species got closer to each other over time, speciation might have been fostered by an increase in competition intensity — another counterintuitive pattern, at least from the niche-filling perspective, as discussed by Drury et al. (2016) in the context of trait evolution under competitive interactions.

This negative correlation between site MNND and speciation in temporal window 3 may reflect competition among ecologically similar species driving character displacement (Brown and Wilson 1956; Schluter 2002). Such competition may mitigate local pressures, resulting in anagenetic speciation at local scale, which would manifest itself at the regional level as budding speciation (Wagner P.J. 1998; Funk and Omland 2003; Otero et al. 2022). Given that those species eventually migrate to other sites, this dynamic aligns with the negative relationship between site diversity and speciation, suggesting a “geographic wave” in which local saturation from a growing regional species pool limits further speciation. Future studies could provide greater insights by investigating the geographic distribution and niche occupation of these species.

Window 4 reveals a contrasting scenario where competition inhibits speciation at the site, as suggested by a positive association with MNND and a negative association with site diversity. By this time, the “cat-gap” was already filled, and ecological opportunities became exhausted. Regional MNND negatively correlates with speciation, diverging from our competition expectations (Sepkoski 1996). Concurrently, regional diversity declines sharply as extinction rates rise (Figures 1C and 4B), suggesting selective extinction targeting species in specific environments (e.g., forests or grasslands) or characteristics. Such selectivity might have removed ecologically similar species that rarely interact locally, increasing regional MNND and creating morphospace gaps. These gaps align with the “species factories” concept, not merely as zones of high turnover, but as potential arenas for the emergence of ecologically distinct and evolutionarily successful species. As environmental instability selectively removes similar taxa, it may open morphospace niches that facilitate the origination of species with broader ecological tolerances and longer durations — consistent with the patterns described by Toivonen et al. (2022).

Environmental changes during the late Miocene further support this process. The transition from browser to grazer-dominated faunas reflects grassland expansion, forest decline, and altered ecological interactions (Strömberg 2002; Figueirido et al. 2015). Cooling temperatures, glacial cycles, and fluctuating habitats exacerbated extinction risks, driving geographic diversity loss. The negative correlation between site diversity and extinction rates indicates that worsening conditions reduced local coexistence. Over time, site diversity declined as extinction increased, supporting a “worst world scenario” where habitat changes and abiotic pressures disproportionately affected Carnivora in North America. These findings highlight how selective extinction, and environmental instability might have shaped diversity at both regional and site levels.

### Eurasia

In Eurasia, early diversification patterns reveal distinct dynamics. During temporal window 1, the arrival of new clades such as Amphicyonidae and Ursidae (Pires et al. 2015) likely contributed to a temporary increase in speciation and extinction rates. This influx of lineages in the Old World likely drove diversification not only among the newcomers but also among pre-existing taxa. It is possible that Amphicyonidae and Ursidae brought a set of novel traits from North America, expanding the morphospace available in Eurasia.

This expansion of morphospace and the resulting diversification can be observed in our competition metric. Regional MNND reveals that as the regional pool became saturated with morphologically similar species, speciation decreased, consistent with competition theory. Over time, declining MNND reflects reduced distances as niches were filled. At the site level, however, MNND shows a negative correlation with speciation alongside a temporal trend of increasing distances (Figure 4E), likely driven by the arrival of ecologically distinct Caniformia immigrants from North America (Pires et al. 2015), and the ecological diversification that started to take place in Eurasia after the arrival of some new ecological forms. These arrivals, and perhaps subsequent *in situ* diversification, expanded the local morphospace, aligning with the ecological opportunity hypothesis, where distinct immigrants are more likely to establish in a new area and perhaps later undergo evolutionary radiations (Mahler and Losos 2010; Yoder et al. 2010).

Environmental conditions during this period also played a critical role, with temperature emerging as a key factor influencing extinction rates. Cooling trends significantly increased extinction risks, as temperature exhibited the most negative exponent among variables. However, higher diversity appeared to buffer some of these effects. This dynamic likely encompasses the effects of what is known as the “Grande Coupure” (∼33.9 Ma), when abrupt climate cooling, habitat shifts, and the arrival of immigrant taxa drastically reshaped faunal communities (Costa et al. 2011). Similar cooling events and habitat changes likely intensified extinction pressures, further influencing the diversification dynamics of early Eurasian Carnivora. It is also worth mentioning that in our dataset Carnivora only existed for a short amount of time during the first time window, and the overall extinction rate shows no drastic changes. Hence caution might be necessary when interpreting all those significant associations between the different metrics and extinction during time window 1.

Non-trivial biotic processes dominated during temporal window 2, as evidenced by the negative relationship between site MNND and speciation. The site MNND time-series is relatively stable for the most part but showed a significant drop towards the end of this time window, which could reflect the potential of interspecific competition to spur speciation rate. Given the correlational aspect of our analysis inferring the direction of causality is not possible, hence such negative association might also simply reflect that new species are more likely to originate very closely in morphospace from their “progenitor” species, even if originally driven by intra-specific competition. Similarly, we find a negative association between reginal diversity and extinction rate. Considering the gradual decline in regional diversity during this time window, this might again indicate selective extinction and that local assemblages start to become more homogenous with morphologically similar species, particularly at the end of the time window.

By temporal window 3, Eurasia experienced a return to relatively long phase of positive diversification, followed by a similarly long period of equilibrium. As observed by Pires et al. (2017a) at the regional scale, speciation rates decrease with rising diversity, although in our case this signal is only recovered at the site scale. This reinforces the idea that diversity-dependence process might be particularly relevant at affecting speciation regimes of Carnivora lineages (Pires et al. 2017a). The morphospace gaps created during the negative diversification that predominated during window 2, combined with improved environmental conditions, likely facilitated this recovery. The onset of the Miocene (∼23 Ma) and the Miocene Climatic Optimum (∼17–15 Ma) provided warm, stable climates that promoted diversification across mammalian clades, including Carnivora (Janis 1993; Koepfli et al. 2008). Grassland expansion during this period created new ecological niches (Janis 1993; Wang and Tedford 2008), driving adaptive radiations and increased speciation. Similar trends are observed in other clades during post-crisis phases, where reduced diversity and morphospace gaps enabled radiation (Foote 2000).

In temporal window 4, these dynamics shifted as diversity-dependent processes began to play a more prominent role in Eurasia. Regional richness increased, intensifying competition and driving higher extinction rates, consistent with a niche saturation model of interspecific competition. Interestingly, site diversity showed a negative correlation with extinction, suggesting that an increase in localized richness might have provided a buffering effect against extinction pressures or that a drop in diversity is associated with further extinction events. Such correlation could indicate that a third factor (external environment) might be driving the fate of diversity at the site level. When diversity drops, potentially driven by an external factor, extinction rises and vice-versa. It is interesting to note that the standard deviation in the number of coexisting species in that time interval (Figure S9 panel B), follows a similar trend as the average number of coexisting species (Figure 4C), with an initial drop, followed by a rise and then a steeper drop towards the present. The standard deviation is also quite small, suggesting that most species coexistence with a relatively similar number of species at the site level, at least when compared to time window 3. This might suggest that the external factor affected local assemblages in a similar manner throughout all Eurasia.

One critical factor that may have influenced these dynamics in Eurasia during the last 7 Ma may be the Messinian Salinity Crisis (MSC) (∼5.96–5.33 Ma), as troughs in regional and site richness around ∼5 Ma align temporally with the event (Hsü et al. 1977; Domingo et al. 2014; Agiadi et al. 2024). Extinction events during the MSC created morphospace gaps, increasing MNND and reducing competition-driven diversification. This is supported by the negative correlation between regional MNND and speciation, and the positive correlation between MNND and extinction, suggesting selective extinction under deteriorating environmental conditions. If certain environments in Eurasia became particularly unfavorable, species reliant on those habitats may have disappeared at a higher rate, effectively reducing local and regional diversity while increasing regional extinction rates. Over time, as fewer species occupied the regional pool, extinction rates may have decreased proportionally, resulting in an eventual stabilization.

### Conclusion

Here we demonstrate that in both North America and Eurasia, competition proves to be a crucial factor influencing diversification. While it can hinder speciation by saturating ecological niches, competition may also promote diversity through processes like character displacement or ecological speciation. We also highlight that we can detect distinct mechanisms act in shaping diversity when measuring competition at local or regional levels, leading to opposite associations among evolutionary rates and our time-series. This complexity mirrors the arguments of Cadotte and Tucker (2017), who discuss how environmental and competitive processes intertwine, producing overlapping yet distinct patterns that vary across spatial and ecological scales. Finally, our findings emphasize the selective nature of extinction. Abiotic forces, such as cooling temperatures or habitat shifts, appear to have disproportionately affected certain environments, selectively removing species and creating “gaps” within the morphospace, which underscores the complexity of diversification dynamics in Carnivora, shaped by an interplay of competition, environmental pressures, and spatial scales. While competition often governs local speciation processes, abiotic factors, particularly selective extinction, emerge as critical drivers at broader scales. By integrating temporal and spatial perspectives, our work provides a deeper understanding of the processes driving macroevolution in Carnivora.

## Declaration section

### Ethics approval and consent to participate

Not applicable.

### Consent for publication

Not applicable.

### Availability of data and material

All data generated or analyzed during this study are included in this published article.

### Competing interests

We have no competing interests.

### Funding

L.M.V.P and T.B.Q. are supported by Fundação de Amparo à Pesquisa do Estado de São Paulo (FAPESP).

### Authors’ contributions

L.M.V.P. and T.B.Q. conceived and designed the study, L.M.V.P. performed the analyses, L.M.V.P. and T.B.Q. wrote the draft.

## SUPPORTING INFORMATION

### Table List

**Table S1.** – The file is available in the appendix of this article.

**Table S2.** – The file is available in the appendix of this article.

**Table S3.** – The file is available in the appendix of this article.

**Table S4.** Number of species and occurrences by taxonomic group and geographic region. The main analyses here were performed by grouping all occurrences from North America and Eurasia into two different pools of species.

| Clades | Species | Occurrence |
| --- | --- | --- |
| Eurasia | 514 | 3702 |
| N. America | 412 | 3647 |
| Caniformia (Eurasia) | 334 | 2102 |
| Caniformia (N. America) | 347 | 3111 |
| Feliformia (Eurasia) | 180 | 1600 |
| Feliformia (N. America) | 65 | 536 |
| Ailuridae (Eurasia) | 19 | 52 |
| Ailuridae (N. America) | 6 | 7 |
| Amphicyonodontinae (Eurasia) | 6 | 20 |
| Amphicyonodontinae (N. America) | 8 | 32 |
| Amphicyonidae (Eurasia) | 35 | 247 |
| Amphicyonidae (N. America) | 40 | 211 |
| Barbourofelidae (Eurasia) | 3 | 24 |
| Barbourofelidae (N. America) | 4 | 33 |
| Canidae (Eurasia) | 38 | 476 |
| Canidae (N. America) | 132 | 1735 |
| Felidae (Eurasia) | 57 | 666 |
| Felidae (N. America) | 42 | 393 |
| Viverridae (Eurasia) | 36 | 94 |
| Viverridae (N. America) | 0 | 0 |
| Herpestidae (Eurasia) | 16 | 34 |
| Herpestidae (N. America) | 0 | 0 |
| Hyaenidae (Eurasia) | 45 | 663 |
| Hyaenidae (N. America) | 5 | 26 |
| Mephitidae (Eurasia) | 15 | 32 |
| Mephitidae (N. America) | 18 | 153 |
| Mustelidae (Eurasia) | 156 | 646 |
| Mustelidae (N. America) | 97 | 566 |
| Nimravidae (Eurasia) | 6 | 48 |
| Nimravidae (N. America) | 11 | 69 |
| Palaeogalidae (Eurasia) | 4 | 16 |
| Palaeogalidae (N. America) | 3 | 15 |
| Percrocutidae (Eurasia) | 9 | 49 |
| Percrocutidae (N. America) | 0 | 0 |
| Procyonidae (Eurasia) | 13 | 32 |
| Procyonidae (N. America) | 28 | 146 |
| Stenoplesictidae (Eurasia) | 4 | 6 |
| Stenoplesictidae (N. America) | 0 | 0 |
| Ursidae (Eurasia) | 52 | 597 |
| Ursidae (N. America) | 18 | 261 |

**Table S5.** – The file is available in the appendix of this article.

### Supplementary Figure List

**Figure S1.**
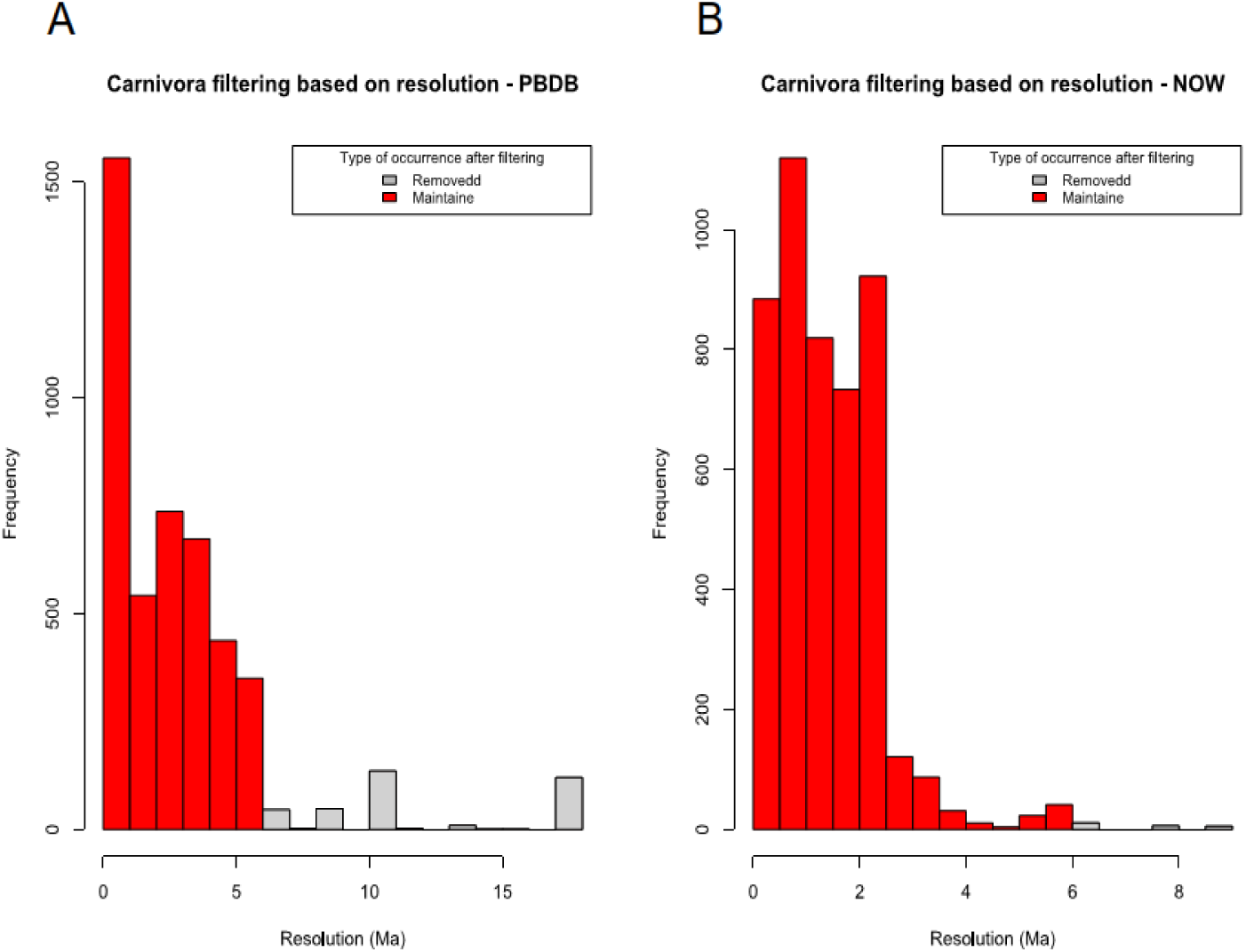
Histogram depicting the resolution of occurrence data from PBDB (A) and NOW (B) datasets. Red bars indicate occurrences maintained in our analyses as they have time intervals lower than 6 Ma. Grey bars represent occurrences that were removed for not attending such a criteria.

**Figure S2.**
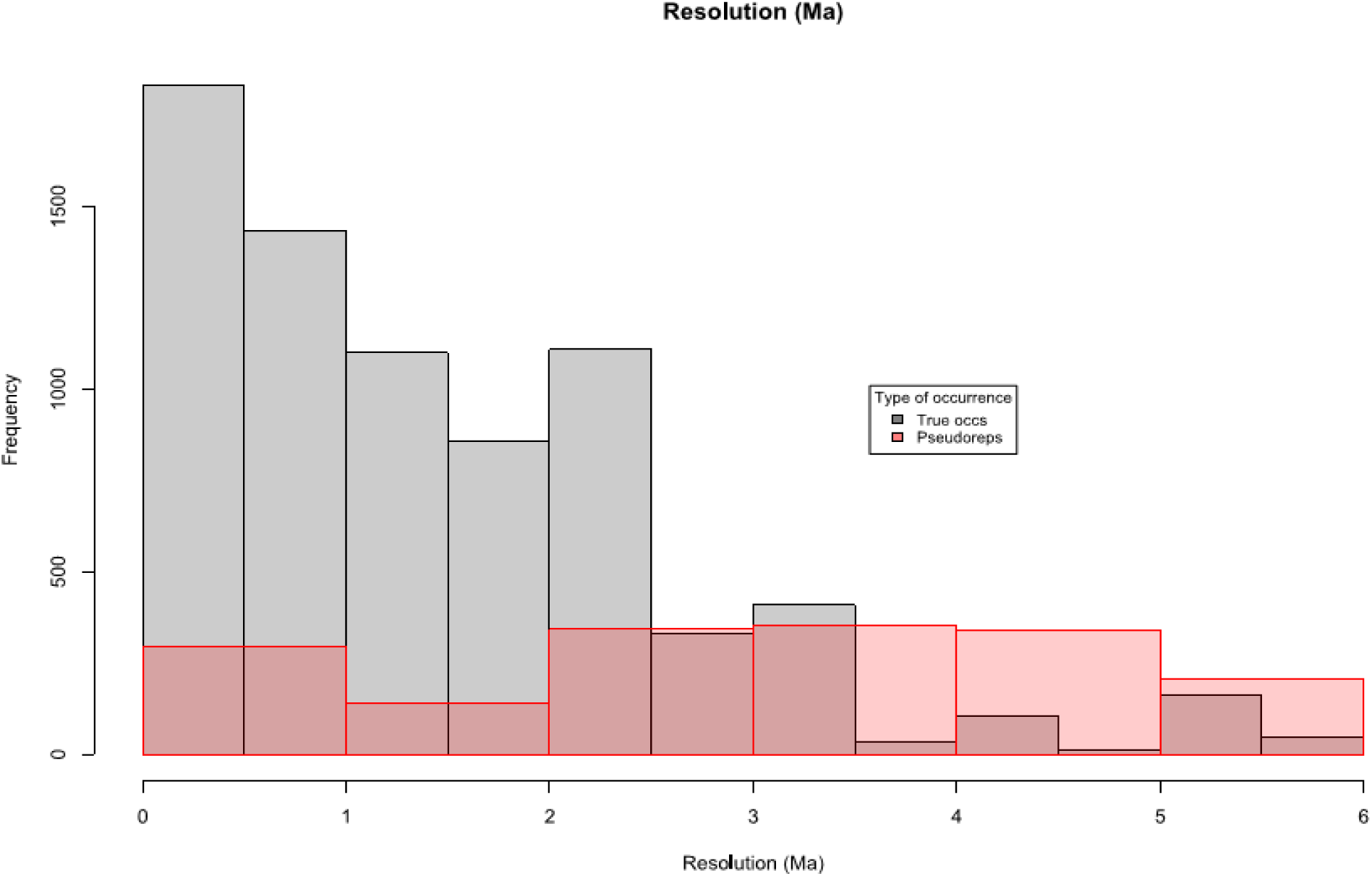
Histogram showing the frequency of pseudoreplicates (potentially replicated occurrences between datasets) among the total number of occurrences in our dataset. All these pseudoreplicates were removed before our main analyses.

**Figure S3.**
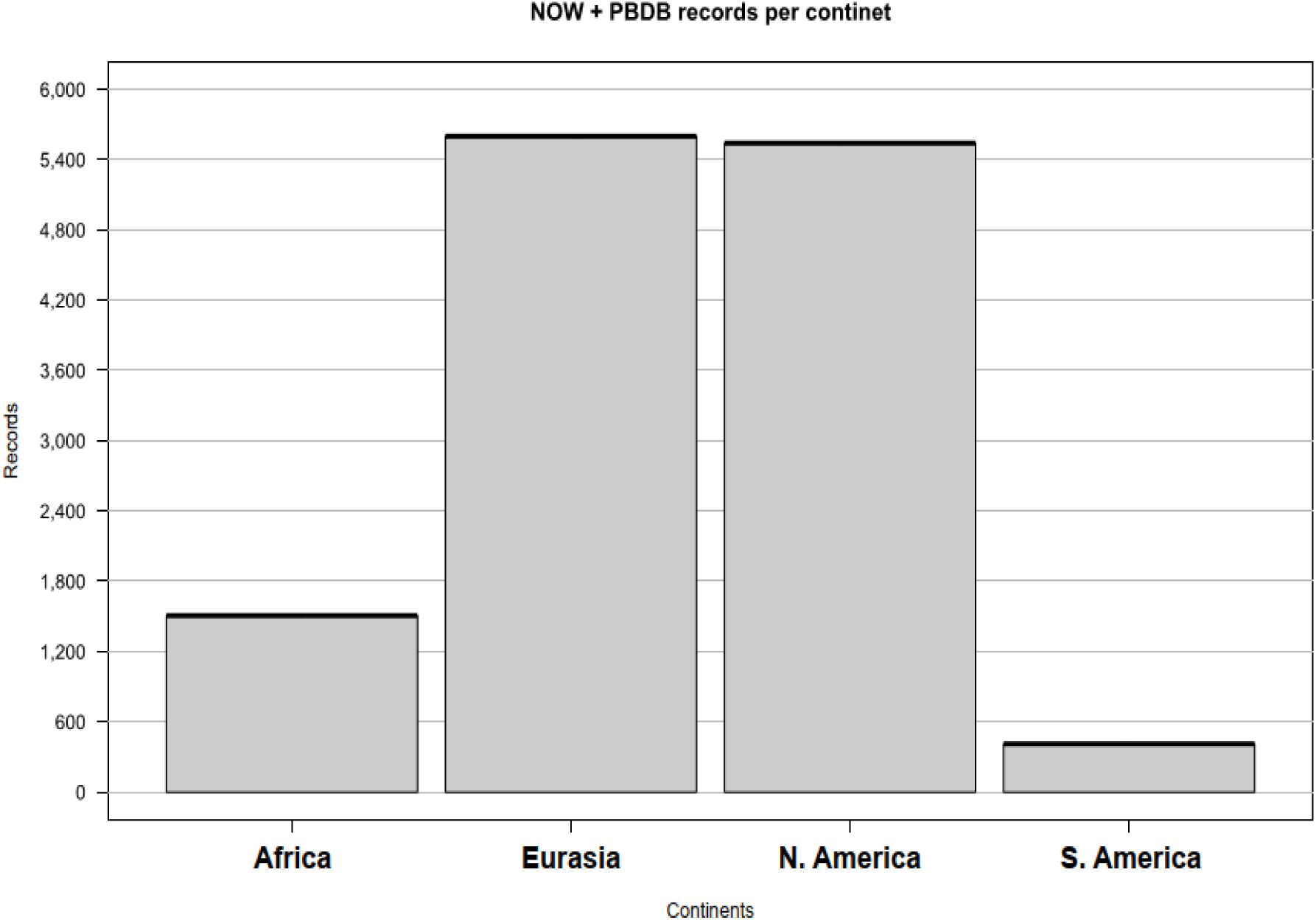
Histogram illustrating the number of occurrences in each one of the four main geographic regions. Such discrepancy that Africa and S. America have compared to Eurasia and N. America was the main reason to exclude both continents in our study.

**Figure S4.**
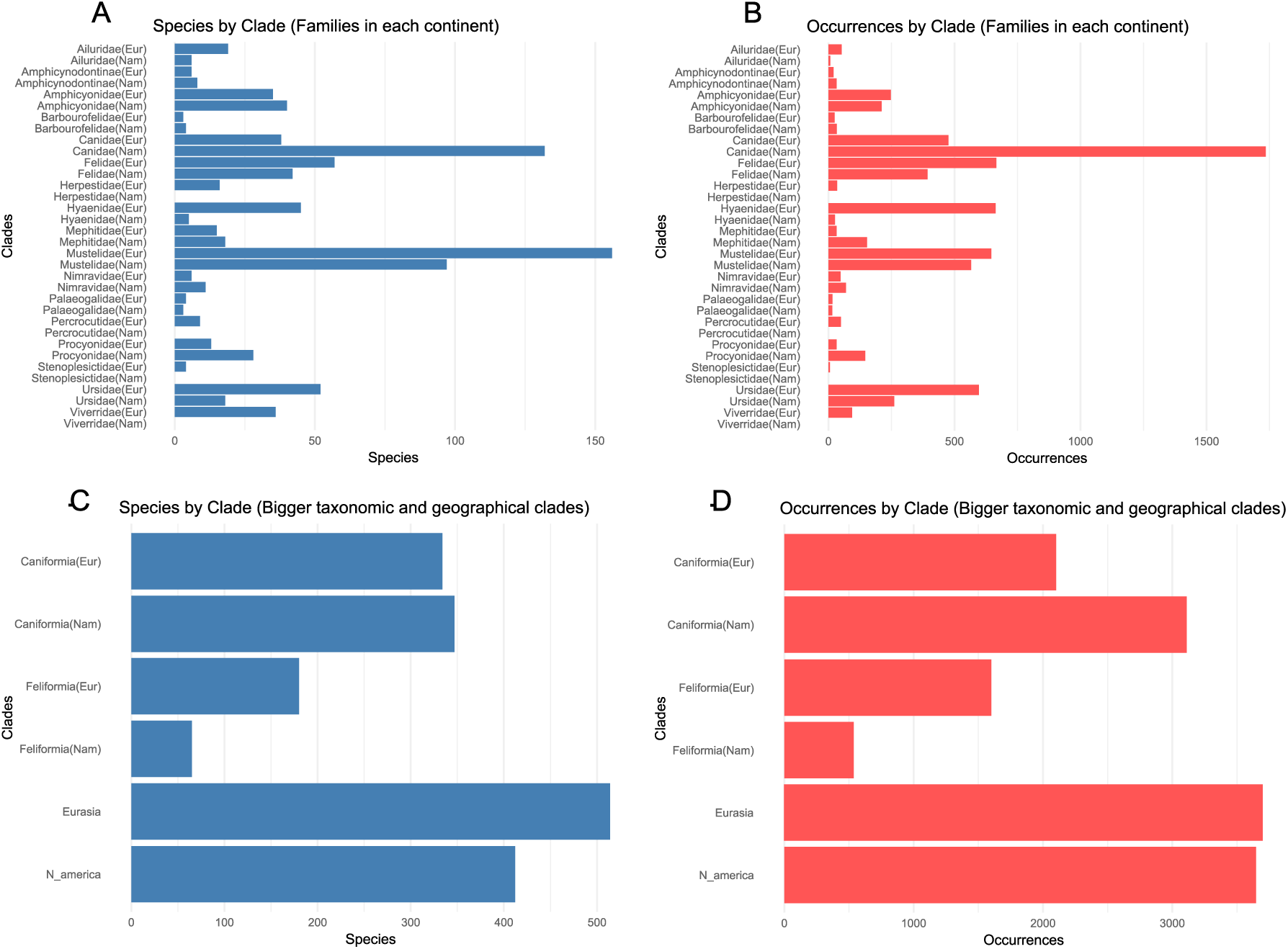
Comparison of species richness (blue) and occurrence records (red) across families and greater taxonomic or geographic groups: (A) Number of species in each family, categorized by geographic region. (B) Number of occurrence records in each family, categorized by geographic region. (C) Number of species in the major taxonomic and geographic clades considered in this study. (D) Number of occurrence records in the major taxonomic and geographic clades considered in this study.

**Figure S5.**
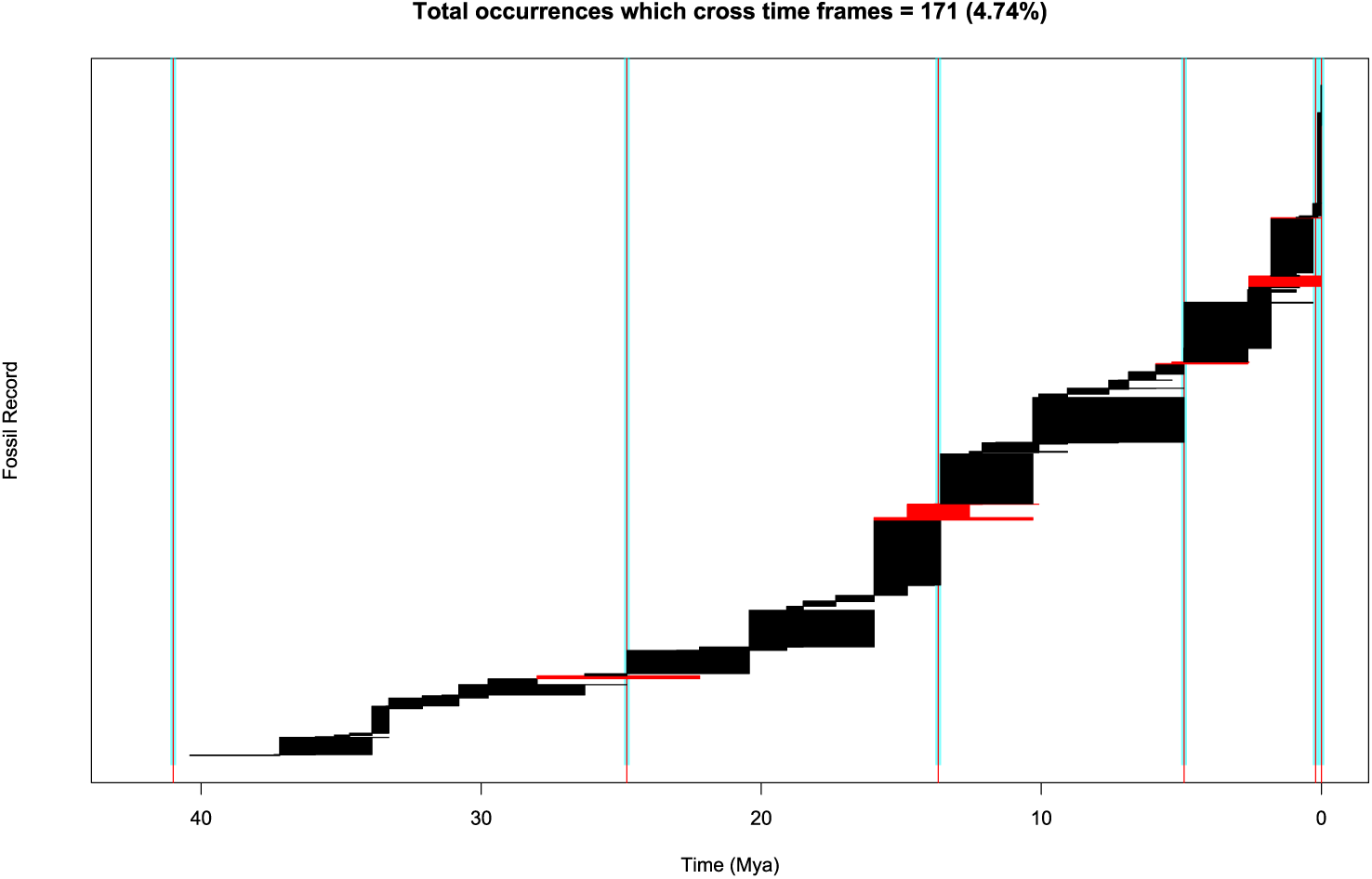
Occurrence temporal distribution and fit to different time windows in North America. Vertical lines correspond to the grouped temporal boundaries at 0, 0.2, 4.9, 13.68, 24.8 and 41 million years. Black bars represent occurrences whose range fits within the intervals described by the boundaries. Red horizontal bars are occurrences whose ranges do not fit any of the grouped time intervals.

**Figure S6.**
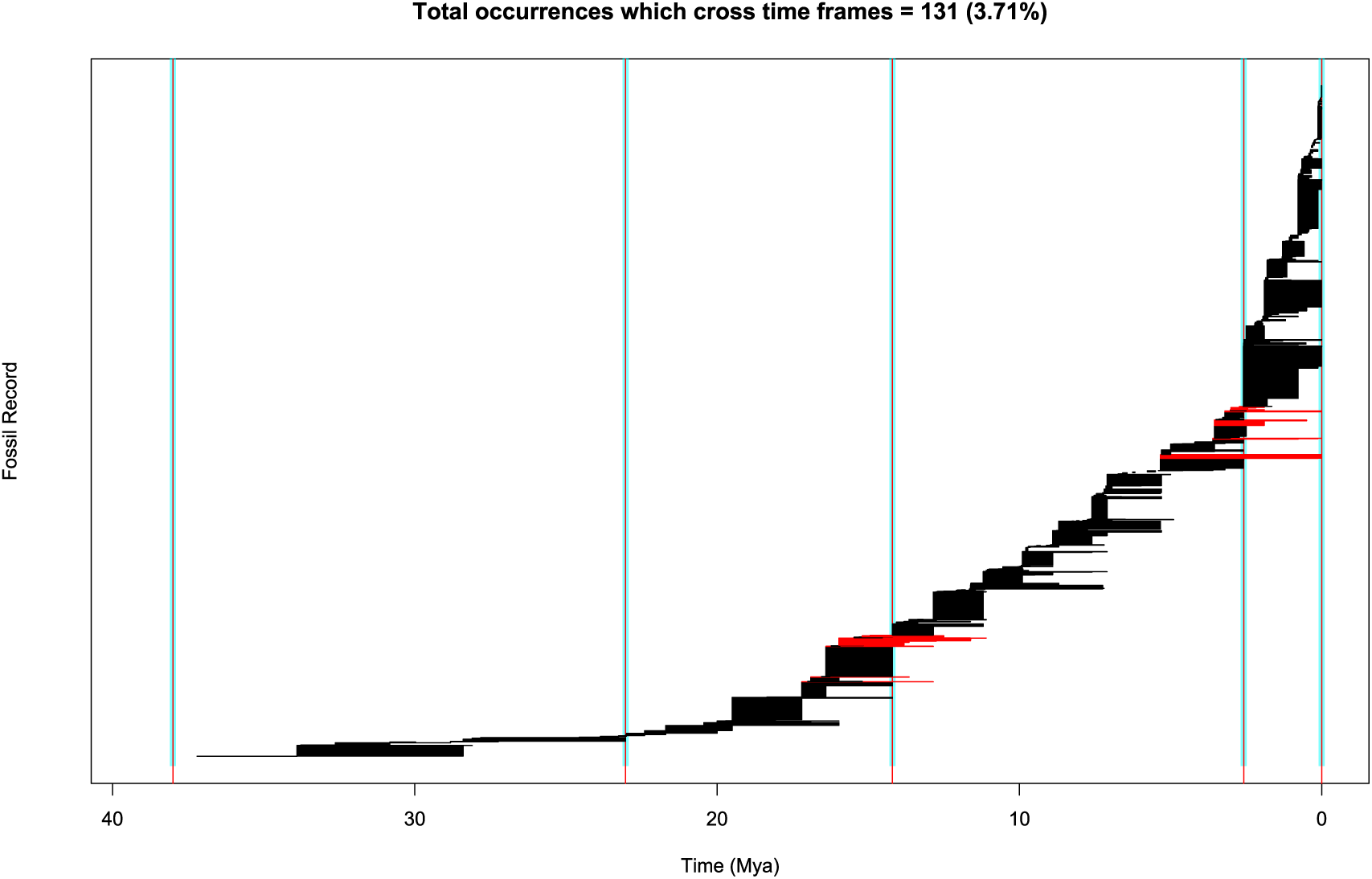
Occurrence temporal distribution and fit to different time windows in Eurasia. Vertical lines correspond to the grouped temporal boundaries at 0, 2.58, 14.2, 23.03 and 38 million years. Black bars represent occurrences whose range fits within the intervals described by the boundaries. Red horizontal bars are occurrences whose ranges do not fit any of the grouped time intervals.

**Figure S7.**
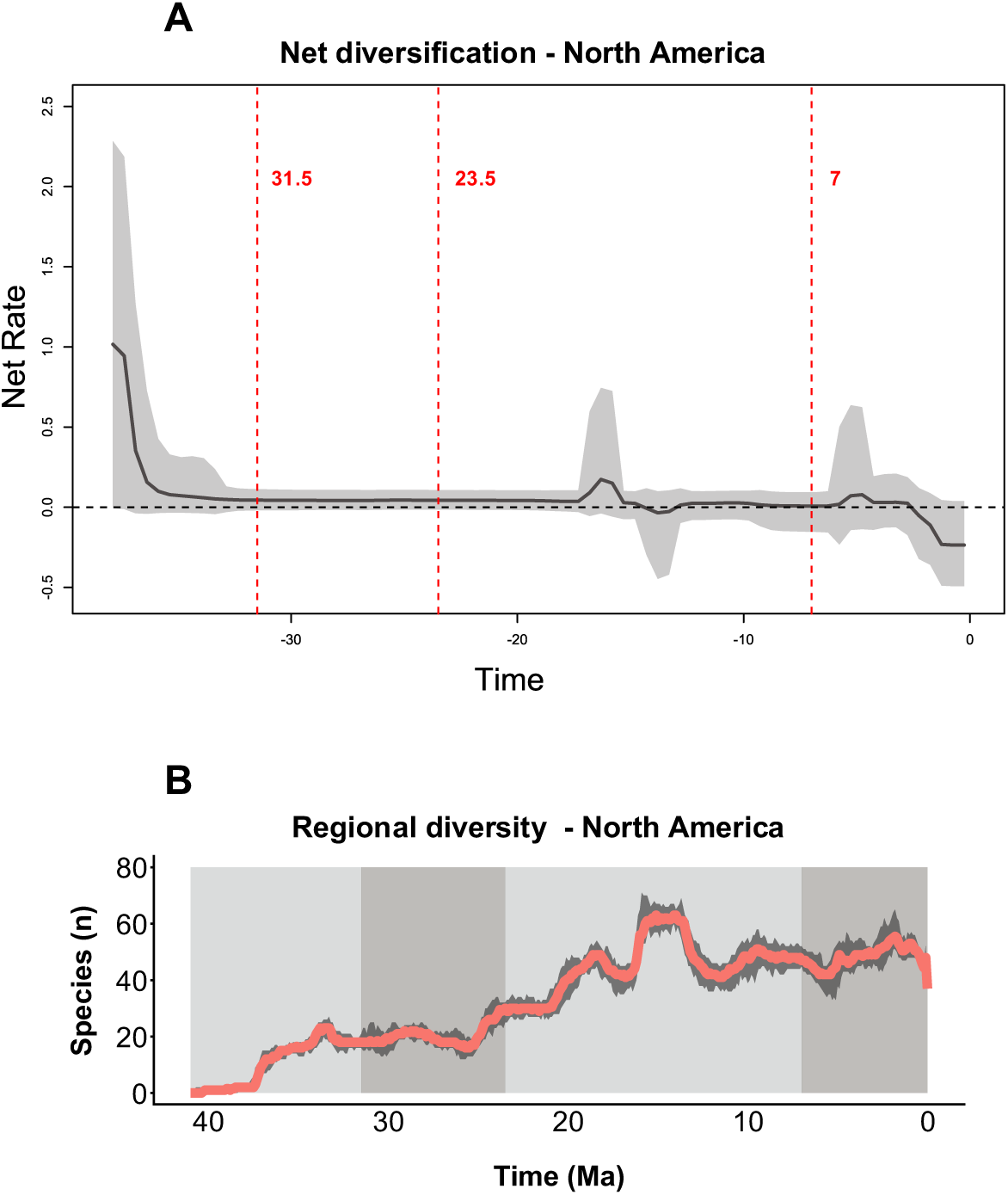
Temporal framework for PyRate MBD analyses using four time windows: (A) Diversification rate of Carnivora in North America over time, highlighting the temporal windows selected for analysis (red dotted vertical lines). (B) Diversity of Carnivora in North America over time, with the temporal windows used in the study represented by distinct shades of gray. The defined time intervals are: window 1: 35–31.5 Ma; window 2: 31.5–23.5 Ma; window 3: 23.5– 7 Ma; window 4: 7 Ma–present. Such intervals were designed to capture periods of radiation, equilibrium, and decline of the clade. These intervals also align with the temporal windows used for analyzing lineages in Eurasia.

**Figure S8.**
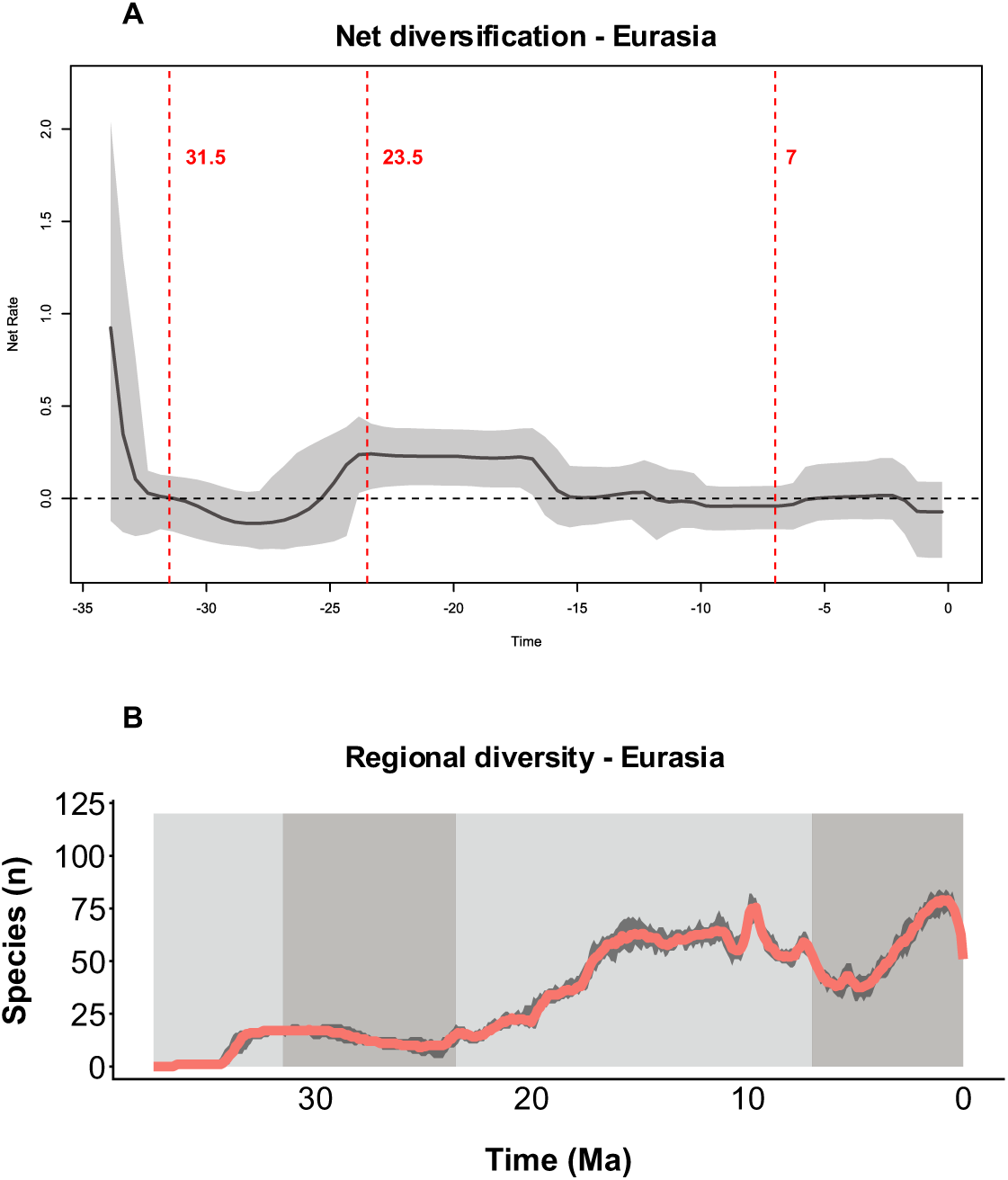
Temporal framework for PyRate MBD analyses using four time windows: (A) Diversification rate of Carnivora in Eurasia over time, highlighting the temporal windows selected for analysis (red dotted vertical lines). (B) Diversity of Carnivora in Eurasia over time, with the temporal windows used in the study represented by distinct shades of gray. The defined time intervals are: window 1: 35–31.5 Ma; window 2: 31.5–23.5 Ma; window 3: 23.5–7 Ma; window 4: 7 Ma–present. Such intervals were designed to capture periods of radiation, equilibrium, and decline of the clade. These intervals also align with the temporal windows used for analyzing lineages in North America.

**Figure S9.**
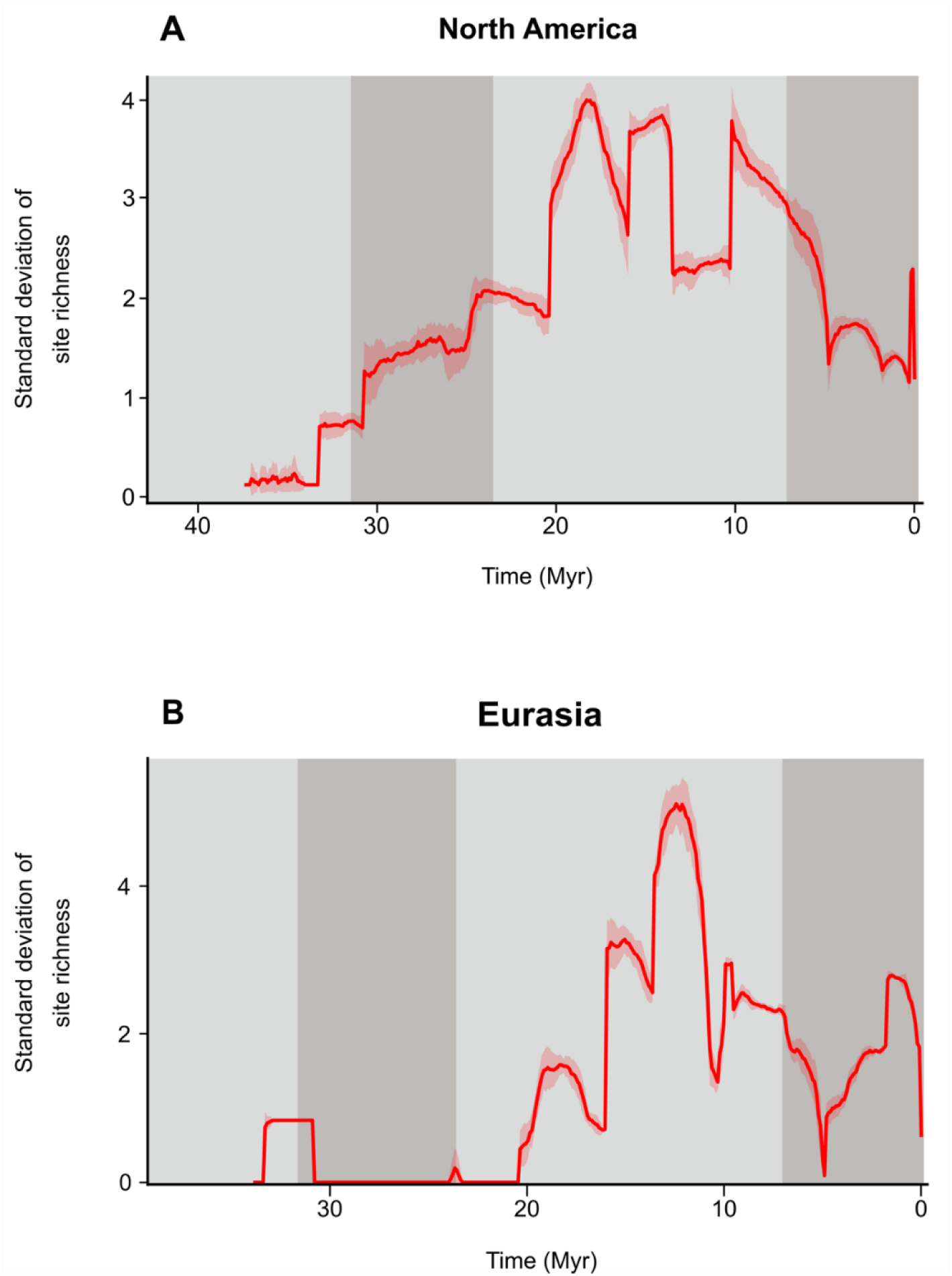
Temporal standard deviation in the number of co-occurring species across 20 replicates for (A) North America and (B) Eurasia. Solid red lines represent the median values, while the shaded areas show the range (maximum and minimum values) across different replicates.

## Notes

### Competing Interest Statement

The authors have declared no competing interest.

